# Fractionation of neural reward processing into independent components by novel decoding principle

**DOI:** 10.1101/2021.08.25.457728

**Authors:** Shitong Xiang, Tianye Jia, Chao Xie, Zhichao Zhu, Wei Cheng, Gunter Schumann, Trevor W. Robbins, Jianfeng Feng

## Abstract

How to retrieve latent neurobehavioral processes from complex neurobiological signals is an important yet unresolved challenge. Here, we develop a novel approach, orthogonal-Decoding multi-Cognitive Processes (DeCoP), to reveal underlying latent neurobehavioral processing and show that its performance is superior to traditional non-orthogonal decoding in terms of both false inference and robustness. Processing value and salience information are two fundamental but mutually confounded pathways of reward reinforcement essential for decision making. During reward/punishment anticipation, we applied DeCoP to decode brain-wide responses into spatially overlapping, yet functionally independent, evaluation and readiness processes, which are modulated differentially by meso-limbic vs nigro-striatal dopamine systems. Using DeCoP, we further demonstrated that most brain regions only encoded abstract information but not the exact input, except for dACC and insula. Furthermore, our novel analytical principle could be applied generally to decode multiple latent neurobehavioral processes and thus advance both the design and hypothesis testing for cognitive tasks.

## Introduction

The brain frequently engages parallel processing involving different latent behavioral processes mediated by functionally distinct, though spatially overlapping, neural networks [1]. Previously, human functional neuroimaging studies have had difficulty in unravelling these processes from basal compound physiological signals [2–4], which has made it difficult to build process-specific and mechanistic models of the brain [5].

Reward/punishment processing is perhaps the most adaptive function of the behavioral control system, optimizing outcomes through both positive and negative reinforcement [6]. Recent overarching frameworks propose two different cognitive processes engaged in parallel during reward or punishment behavior, namely evaluation (i.e., scaling signal values from reward to punishment) and response readiness (subsuming arousal and attentional salience, contributing to response preparatory processes) [6, 7]. The evaluation process is essential for guiding upcoming action selections based on their value, for which the brain has evolved dedicated regions/circuits [8–13]. Complementary to evaluation, both reward and punishment, as highly salient events, attract greater attention than neutral stimuli, also engaging greater levels of motor preparation and emotional arousal [14–16], hence contributing to response readiness. Therefore, evaluation and readiness signals are inevitably confounded with each other during reward/punishment processing. Unfortunately, decomposing this compound signal, for example in human fMRI studies, has proven challenging because these two components cannot be identified by using only reward (or only punishment) stimuli in many experimental paradigms. Previous attempts have been made to overcome this problem by decoding evaluation and readiness signals using the trial-level or first-level data and identifying their corresponding spatially dissociated brain regions [2, 3, 17]. However, these approaches have failed to disentangle signals in brain regions known to encode both evaluation and readiness signals, for example, in the striatum and the ventromedial prefrontal cortex (vmPFC) [7, 16, 18]. Further, the existing studies have not provided convincing evidence to clarify the assumption of functional independence of the evaluation and readiness processes.

To resolve this complex theoretical issue, we developed a novel analytical approach, orthogonal-Decoding multi-Cognitive Processes (DeCoP). Through this innovative approach, for the first time, we have achieved a brain-wide voxel-wise orthogonal decomposition of process-specific neural representations of complex neurobehavioral processes. We further demonstrated that our approach not only provided a valid theoretical statistical inference for the functional independence between the spatially overlapping signals but had wide application for decoding the latent neurobehavioral processes from compound neuroimaging signals, hence further advancing our understanding of the neural basis involved in cognitive processing.

## Results

### Participants and experiment designs

A modified version of the monetary incentive delay (MID) task [19], one of the classical and widely used fMRI paradigms for reward processing, was conducted in 1939 children aged 9-11 from the Adolescent Brain Cognitive Development (ABCD) cohort [20] (Table S1). The MID task consists of five levels of incentive: large loss, small loss, neutral, small win and large win (i.e., −5.0, −0.2, 0, 0.2 and 5.0 $ respectively, Fig. 1a; also see Supplementary Methods for details). There was a discrepancy between the undifferentiated behavioural performance in the contrast of reward vs punishment (Reaction Time: *t_1,927_* = −1.65, *Cohen’s d* = −0.04, *p* = 0.10; Accuracy: *t_1,927_* = 0.71, *Cohen’s d* = 0.02, *p* = 0.48; Fig. 1c) and the unbalanced corresponding activation in two of the most critical brain regions (i.e., the vmPFC: *t_1,927_* = 9.83, *Cohen’s d* = 0.22, *p* <1E-21 and the striatum: *t_1,927_* = 10.77, *Cohen’s d* = 0.25, *p* <1E-25; Fig. 1b), hence indicating that simple contrasts for activation detection may not be sufficient to capture latent neurobehavioural processes underlying tasks with multiple cognitive processes.

**Fig. 1.**
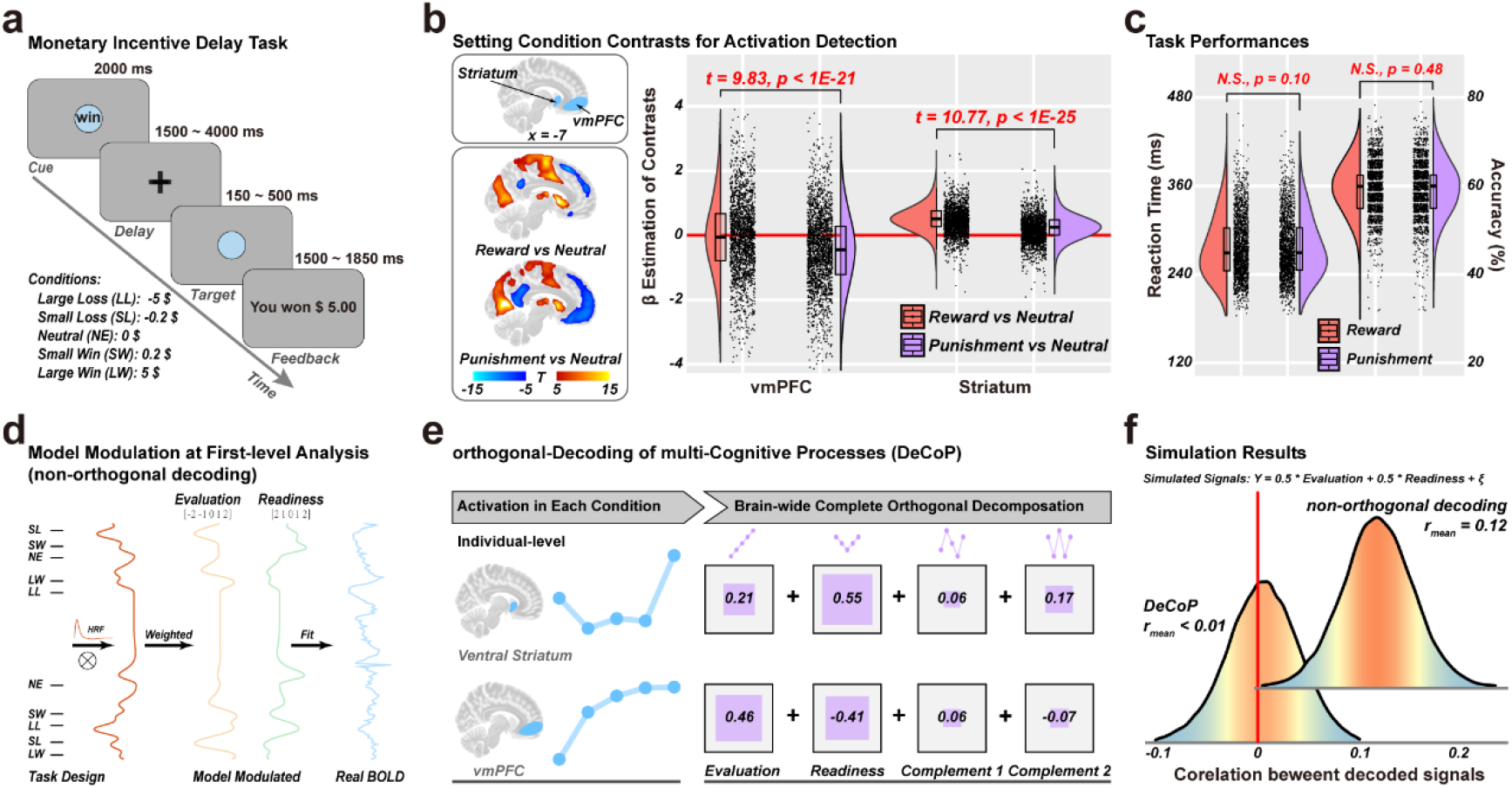
The experimental design and decoding approaches. **a.** Procedure of the monetary incentive delay (MID) task. **b.** Activation detection with setting simple contrasts of task conditions. **c.** The task performances (including reaction time and accuracy) among the participants. **d.** An illustration of univariate decoding at first-level. **e.** An illustration of our novel approach, orthogonal-Decoding of multi-Cognitive Processes (DeCoP). **f.** The simulation results of decomposed signals with univariate decoding and DeCoP. Also, see Methods for details of the simulation.

### A novel orthogonal decoding approach

Previous attempts have tried to decode the latent neural representation of different cognitive processing signals using a model-based linear decomposition at the first level analysis. Briefly, for each individual, each trial (i.e., one of −5.0 $, −0.2 $, 0, 0.2 $ or 5.0 $) was assigned with a corresponding index from either the evaluation model [−2, −1, 0, 1, 2] or the readiness model [2, 1, 0, 1, 2], and then the observed BOLD signals were regressed against either series of trial-wise indices (convolved with the HRF) in a linear model to compute the corresponding univariate model contribution (Fig. 1d, also see Supplementary Methods for details). However, the above approach suffers a significant drawback: while both evaluation and readiness models are designed to be ‘orthogonal’ (i.e., the inner product is null), their actual trial-wise indices used in the actual linear models can hardly be uncorrelated (Fig. S1a upper). This can be attributed to both a systematic bias introduced by the process of convolution with HRF and unbalanced trial numbers that may arise even from a balanced design, e.g. different failing rates across experimental conditions. Using a real data simulation with two independent latent signals (see Methods for details), we demonstrated that the univariate estimates of signal strength could seriously deviate from the simulated value due to the signal admixture (Fig. S1a lower) and decoded signals are inevitably correlated (*r*_mean_ = 0.12, Fig. 1f, also see Supplementary Results for more details). Therefore, the decoding approach based on time series data generally failed to acquire orthogonal signal decompositions (referred to as ‘non-orthogonal’ in the rest of the manuscript), and hence no meaningful inference for the independence of underlying latent processes could be made.

Here, we introduced a novel approach, the orthogonal-Decoding of multi-Cognitive Processes (DeCoP), that not only can provide a model-based unbiased orthogonal decomposition at the condition level, but also enables a statistical evaluation of whether the decomposed signals are indeed independent (also see Methods for details). The central idea to note is that there are five experimental conditions in the MID task that can evoke condition-specific neuronal responses, thus allowing four underlying orthogonal contrasts (or patterns of responses) over the five conditions plus a constant term. Crucially, these orthogonal contrasts should have a clear interpretation, by design, in terms of latent neurobehavioural processes. Specifically, in the second-level analyses of BOLD signals, two readily plausible orthogonal contrasts (i.e., their covariance equals 0) are evaluation (i.e., [−2, −1, 0, 1, 2]) and readiness (i.e., [2, 1, 0, 1, 2]) that respectively reflected putative hypothetical processes of value and salience information across five task conditions. In addition to the two primary contrasts above, their complementary orthogonal contrasts (i.e., the N-shape model: [−1, 2, 0, −2, 1] and the W-shape model: [1, −2, 2, −2, 1]) are also available and can explain information not accounted for by the hypothetical contrasts for evaluation and readiness. With the above orthogonal settings, we were thus able to retrieve the orthogonally decomposed signal components of underlying latent processes and assess their respective representations over the entire brain (Fig. 1e). Notably, DeCoP allows us to make meaningful inference regarding the independence of underlying latent processes, i.e., uncorrelation is equivalent to independence in our settings (real data simulation |*r*_mean_| < 0.01, Fig. 1f, also see Supplementary Methods and Results for the detailed proof).

### Decomposed neural representations with DeCoP

In our initial report of the results, we will focus on the evaluation and readiness components. Notably, the vmPFC (Brodmann area [BA] 10-11; Peak MNI: [−9, 49, −9], Cluster: 615 voxels, *p*_FWE-corr_ = 1.87E-08) and ventral striatum (VS, Peak MNI: [−7, 25, −3], Cluster: 634 voxels, *p*_FWE-corr_ = 1.36E-08) were the most prominent regions identified in the evaluation model (Fig. 2a upper left & 2b), thus being highly sensitive for tracking the entire dimension from punishment to reward. These areas coincide with the terminal regions of the dopamine neuron projections from the ventral tegmental area (VTA), i.e. the meso-corticolimbic dopamine system [21–23]. For the readiness model, however, the signals were more widely dispersed across cortical and subcortical areas, including motor-somatosensory, salience and attention networks, and regions such as the dorsal striatum (DS, Peak MNI: [8, 10, 4], Cluster: 1688 voxels, *p*_FWE-corr_ = 1.75E-14) and thalamus (THA, Peak MNI: [13, −6, 16], Cluster: 2267 voxels, *p*_FWE-corr_ = 1.11E-16) (Fig. 2a upper right & 2b), consistent with their engagement in processing both reward and punishment [24].

**Fig. 2.**
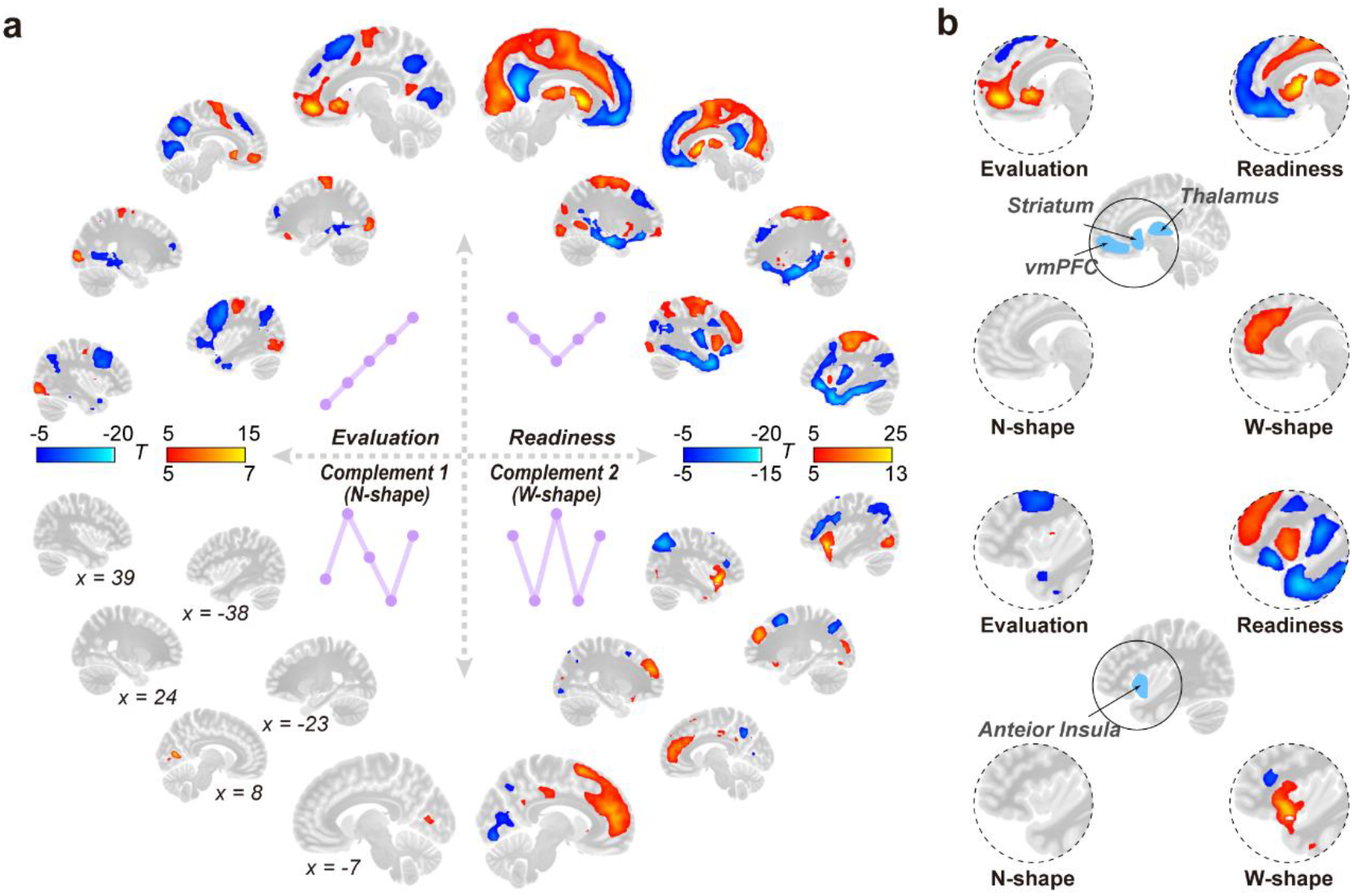
Neural representations of the orthogonal decomposition. **a.** Brain-wide T-maps of decomposed signals for orthogonal contrasts. **b.** Decomposed signals in highlighted brain regions. Brain-wide significance was set as ļTļ > 5, i.e., significant with voxel-wise Bonferroni correction. The MNI coordinates of brain slices were inserted at the lower left.

### Neural circuits for decomposed signals

We then investigated whether the neural representations of evaluation and readiness signals were underpinned by different neural circuits, in particular those modulated putatively by the midbrain dopaminergic projections originating from either the substantia nigra pars compacta (SNc) or the VTA, which plays a central role in reward prediction and approach [21, 22]. We found regions of the evaluation model with higher functional connectivity (FC) to VTA than to SNc (*paired t*-test: *t_183_* = 14.84, *Cohen’s d* = 1.10, *p* < 10E-32), and regions of the readiness model with higher FC to the SNc than to the VTA (*paired t*-test: *t_183_* = 3.63, *Cohen’s d* = 0.27, *p* = 0.0004, Fig. 3a) based on 7T high-resolution resting-state fMRI data from the Human Connectome Project (HCP) [25]. Further, we extracted the *t*-maps of the difference between the seed-based FC from VTA and SNc (i.e., ‘VTA > SNc’) (Fig. 3b), which exhibited high similarities, although in opposite directions, with the *t*-maps of both evaluation (*r* = 0.22, *p_adj_* < 10E-20) and readiness (*r* = −0.12, *p_adj_* < 10E-12, Fig. 3c) models. Thus, the separate VTA and SNc dopamine projections could be the putative source of evaluation and readiness signals, respectively.

**Fig. 3.**
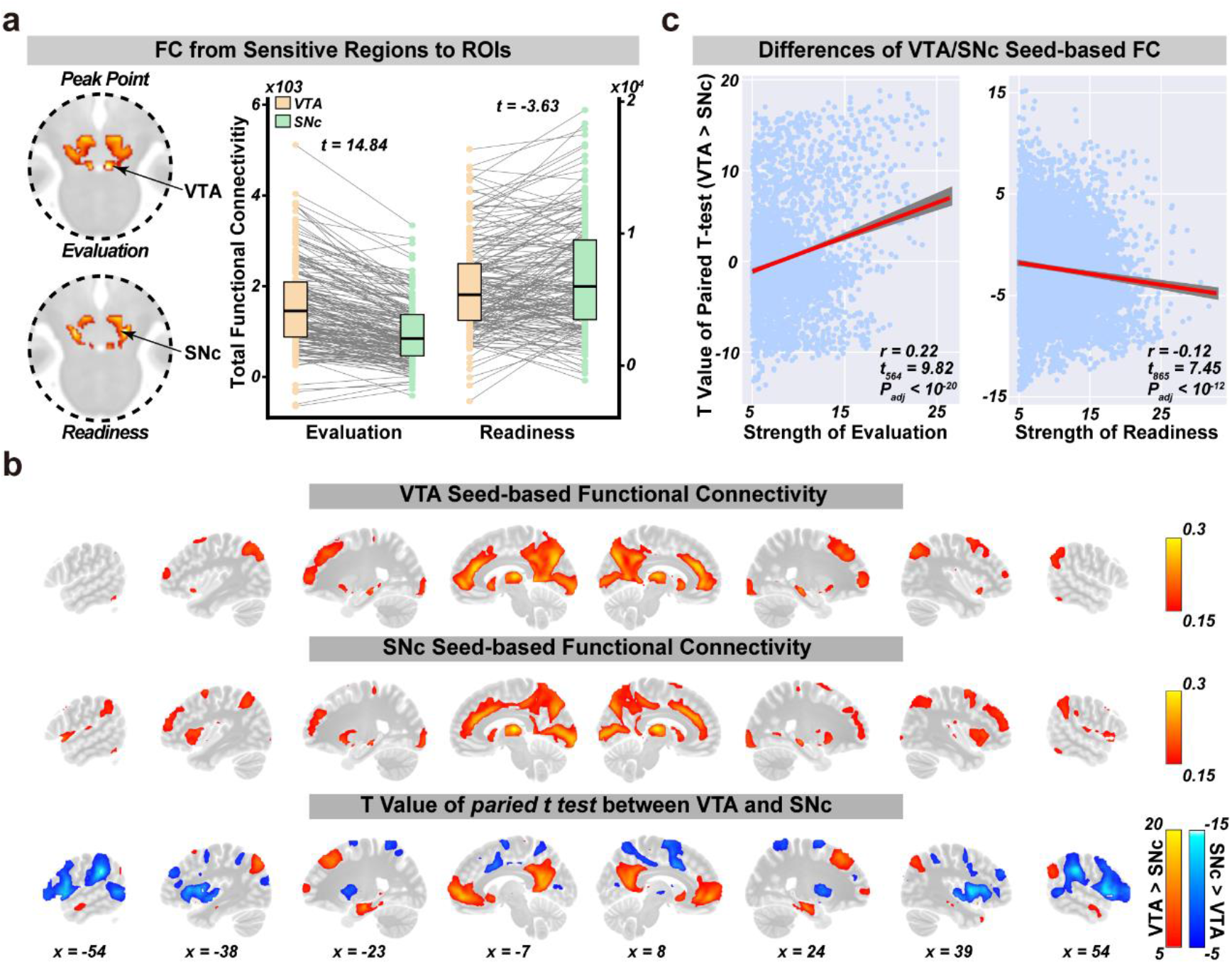
The decomposed evaluation and readiness processing targets VTA and SNc neural circuits, respectively. **a.** Left: strength of functional connectivities (FCs) to VTA and SNc from the evaluation and readiness regions identified in Fig. 2. Right: Paired t-tests between FCs to VTA and SNc from evaluation and readiness. **b.** The brain-wide patterns of seed-based FC from regions of interest (ROIs). Top: VTA seed-based; Middle: SNc seed-based; Bottom: The T-map of the differences between seed-based FCs patterns from VTA and SNc. **c.** Brain-wide pattern similarities between the strength of decomposition signals (left: evaluation; right: readiness) and the differences of seed-based FCs from VTA and SNc.

### Decomposed processes affect distinct cognition components

We further implemented a weighted voxel co-activation network analysis (WVCNA, see Supplementary Methods for details) to capture the most informative brain-wide signal clusters [26] and identified 55 and 194 clusters for the evaluation and readiness processes, respectively (Fig. S2 & Table S2-3). Using the canonical correlation analysis (CCA, see Supplementary Methods for details), we then found significant associations between variations in the decomposed neural signal and distinct cognitive components across eight reward-processing-related behaviours for both evaluation and readiness (evaluation: adjusted η^2^ (*adj-η^2^*) = 0.033, *p_perm_* = 0.0241; readiness: *adj-η^2^* = 0.113, *p_perm_* = 0.0001, Table S4). For the evaluation process, higher behavioral inhibition and crystallized intelligence were mainly associated with reduced sensitivity in the bilateral inferior temporo-occipital junction, middle cingulate cortex, nucleus accumbens (NAcc) and left dorsal anterior cingulate cortex (dACC), and hyperactivity of these regions may lead to internalizing disorders (such as anxiety and depression) (presented by the first component, Fig. 4a upper & Table S5). Further, higher activations in the vmPFC and subgenual ACC (sgACC) were also associated with fun-seeking and externalizing problems (such as rule-breaking and aggressive behavior) (presented by the second component, Fig. 4a lower & Table S5). The readiness process seemingly involved two competing processes regulating the adaptation of external stimulus and incentive salience. Specifically, the first cognition component may represent the ability of positive reinforcement learning and flexibility of adaptation, given the positive loadings of reward responsiveness and fluid intelligence as well as the negative loading of internalizing scores, which were mainly negatively associated with dACC, rostral ACC (rACC), right thalamus, putamen, insula and inferior/middle frontal cortex (IFC/MFC) (Fig. 4b upper & Table S6). Nevertheless, the dACC and right insula, commonly considered the critical regions of the salience network, were also activated by incentive motivation (i.e., higher reward responsiveness and drive in the third component, Fig. 4b lower & Table S6). Differentiated activations between the left and right sensorimotor area combined with positive activations of right IFC/MFC and insula may contribute to the action control, hence leading the persistent pursuit of desired goals [27](presented by the second component, Fig. 4b middle & Table S6).

**Fig. 4.**
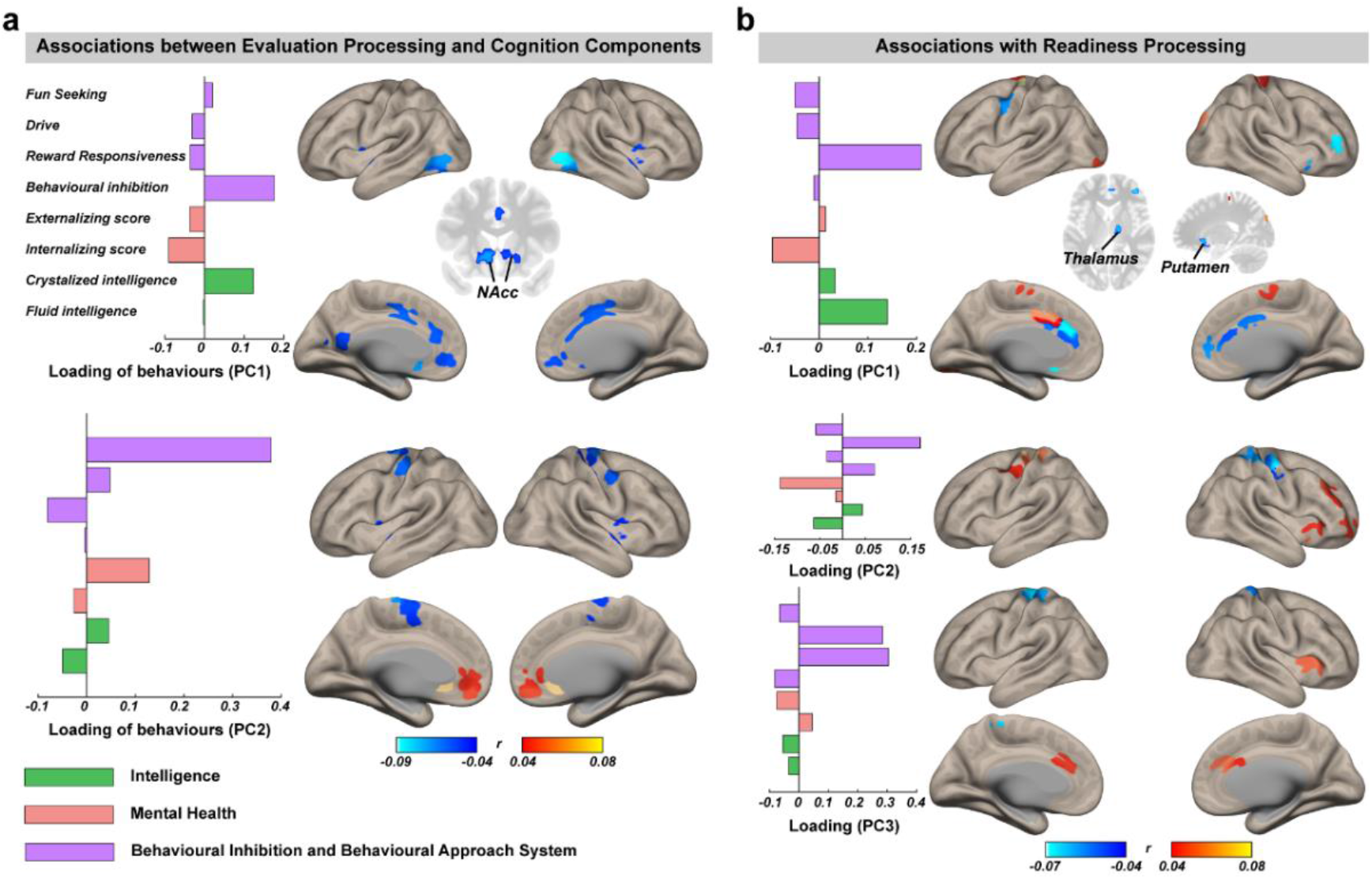
Impacts of evaluation and readiness processing on cognitive function. The canonical correlation analysis (CCA) was implemented to further segregate signal components that demonstrated differential associations with distinct cognition components (**a.** for evaluation processing and **b.** for readiness processing). The loadings of behaviors are shown in the left of the subgraphs. The loadings of brain activations were represented by the correlations with the cognition components. Only the significant regions were reported in the subgraphs. Also see Table S4-6.

### Independence of evaluation and readiness

We further demonstrated that the above spatially overlapping cognitive processes modulated by distinct neural pathways were indeed functionally independent, which could be directly inferred from uncorrelated signal components at the co-activated regions (see Supplementary Methods for the detailed proof). Based on our simulation results, if and only if the compound signals were indeed a combination of independent signals, and the correct orthogonal contrasts were applied, the decomposed signals could be uncorrelated (|*r*_mean_| < 0.001, the ‘Independent’ model). Otherwise, the decomposed signals were highly correlated and hence inseparable as modulations of latent signals (the ‘One Signal’ model: *r*_mean_ = 0.54; the ‘Push and Pull’ model: *r*_mean_ = −0.45; Fig. 5a & Table S7).

**Fig. 5.**
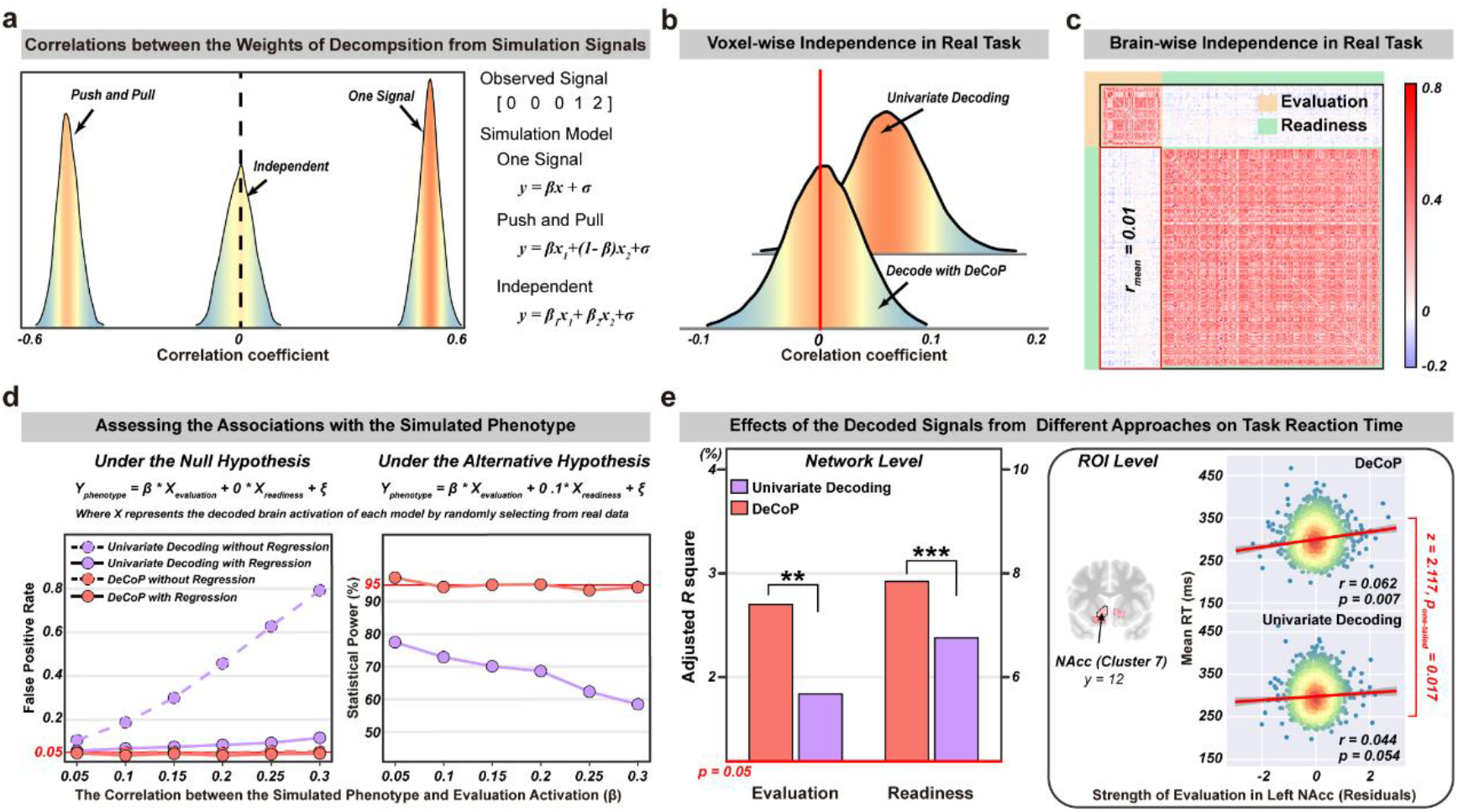
Valid infernece for functional independence and higher statistical power provided by DeCoP. **a.** Correlations between decomposed signals based on different simulation models. Also see Table S7. **b.** The distributions of the correlations between the decomposed signals at each voxel. **c.** The correlation matrix of signals from the evaluation and readiness clusters identified by WVCNA. **d.** With simulated phenotypes, DeCoP provided less false positive rate and higher statistical power. **e.** In the real task, DeCoP also provided more variance explained at both the network level and the ROI level (Left NAcc as an instance). Also see Fig. S3.

Using DeCoP we have thus demonstrated that evaluation and readiness are indeed functionally independent processes at each co-activated voxel across the whole brain (*r_mean_* = 0.006, 95%CIs = [-0.009, 0.021], *p_bootstrap_* = 0.4142, where 99.5% voxels with |*r*| < 0.1; Fig. 3b), while traditional ‘non-orthogonal’ decoding would find both decomposed signals to be significantly correlated (*r*_mean_ = 0.071, 95%CIs = [0.044 to 0.103], *p_bootstrap_* < 0.0001; Fig. 5b). We also observed brain-wide low inter-correlations between evaluation and readiness clusters (*r*_mean_ = 0.01, range = −0.08~0.11, *p_bootstrap_* = 0.8379; Fig. 3c), in contrast to the very high intra-correlations within each of evaluation and readiness clusters (*r*_mean_ = 0.345, *p_bootstrap_* = 0.0042, Fig. 5c), hence further supporting their neural functional independence. Therefore, the observed unbalanced sensitivity towards reward and punishment in brain regions such as VS and vmPFC (Fig. 1b) could be parsed into two spatially overlapped though functionally independent balanced signal components.

### Fewer false inferences using DeCoP

With simulated phenotypes (Fig. 5d, also see Methods for details), we first demonstrated that univariate ‘non-orthogonal’ decoding could lead to a seriously inflated false positive rate because the thus decomposed signals are a mixture of underlying latent components (Fig. 5d Left). An intuitive correction for this signal admixture was to mutually control for the other non-orthogonally decoded component. However, while this mutually control approach could largely alleviate the inflated false positive (Fig. 5d Left), it also significantly reduced the statistical power (Fig. 5d Right), again because of the signal admixture. In contrast, DeCoP could provide uniformly better performance with a properly controlled false positive rate and greater statistical power across all simulation settings (Fig. 5d).

In the real data, the mean reaction time of the MID task could be found in association with both evaluation and readiness processing with either DeCoP or non-orthogonal decoding after mutually controlling for both processes (Fig. 5e & Fig. S3). However, DeCoP demonstrated significantly increased statistical power, i.e. exhibited more explained variance than ‘non-orthogonal’ decoding at both the network level (> 45% additionally explained variance exceeding the significant threshold at 0.05, *p_perm_*< 0.01; Fig. 5e Left & Fig. 3) and the ROI level (for instance, the NAcc could only be identified with DeCoP with twice the variance explained; Steiger’s test *Z* = 2.11, *p_one-tailed_* = 0.02; Fig. 5e Right & Fig. S3). Hence, the functional independence advocated by our novel approach of DeCoP is essential for revealing independent neurobehavioral processes.

### Complementary components using DeCoP

Additionally, we found that the signals attributed to evaluation and N-shape models (dependent signals: *r*_mean_ = −0.093, *p_bootstrap_* < 0.0001) together described the sensitivity of evaluation from punishment to reward. Further, signals of evaluation and N-shape models were independent (evaluation *vs* W-shape: *r_mean_* = −0.008, *p_bootstrap_* = 0.3358; N-shape *vs* readiness: *r_mean_* = −0.001, *p_bootstrap_* = 0.8432) of those attributed to the readiness and W-shape models (dependent signals: *r*_mean_ = −0.159, *p_bootstrap_* < 0.0001), which together described the differentiated engagement of readiness from the neutral condition to reward/punishment conditions (Fig. 6a). Hence, the complementary N-shape and W-shape models account for the deviation from the latent evaluation and readiness signals of the proposed evaluation and readiness models respectively (Fig. 4b). The N-shape model was only observed with significant signals in the primary visual cortex (BA 17; Peak MNI: [4, −81, 1], *t_1,927_* = 7.65, *Cohen’s d* = 0.17, Cluster: 233 voxels, *p*_FWE-corr_ = 2.91E-05, Fig. 2a lower left & 2b). For the W-shape model, the most prominent regions were bilateral anterior insula (aINS, BA 38, Peak MNI: [49, 25, −9], *t_1,927_* = 14.83, *Cohen’s d* = 0.34; Cluster: 1177 voxels, *p*_FWE-corr_ = 4.41E-05) and anterior cingulate cortex (ACC, BA 32, Peak MNI: [7, 49, 22], *t*_1,927_ = 11.51, *Cohen’s d* = 0.26; Cluster: 881 voxels, *p*_FWE-corr_ = 2.87E-10, Fig. 2a lower right & 2b). We further demonstrated that the signal strength of the additional complementary orthogonal contrasts could provide a useful measurement of the distance between the latent independent signals and the proposed models (see Supplementary Methods for details). Converging evidence indicated that most brain regions distinguish reward from punishment signals with their relative rank, hence processing highly abstract information only (Fig. 5c-d & Table S8). However, the bilateral aINS and dorsal ACC were most likely tracking the parametric nature of the experimental design (i.e., [−5, −0.2, 0, 0.2, 5]; Fig. S4; also see Supplementary Results for more details).

**Fig. 6.**
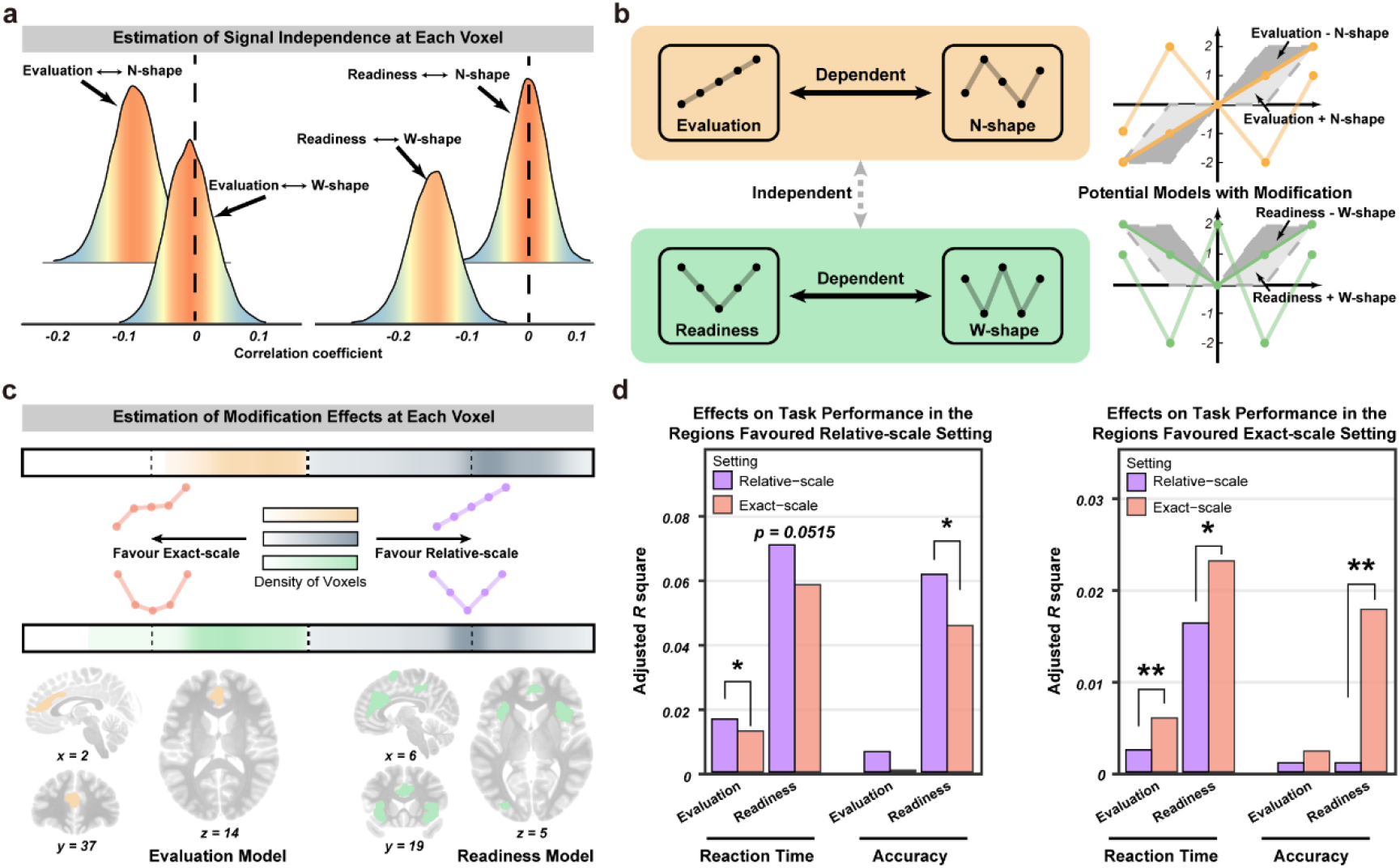
The modulation effects of the additional complementary orthogonal contrasts and the optimal models for latent neurobehavioral signals. **a.** The distributions of pair-wise correlations between signals of orthogonal contrasts at each voxel. Mean correlations deviated from 0 would indicate a pair of related signals. **b.** An illustration of how related signals could describe the evaluation-related and readiness-related processing. **c.** The bimodal distribution of voxels favoring the exact-scale or relative-scale settings in evaluation and readiness processing. The typical regions favoring the exact-scale setting were illustrated in the corresponding lower subplots. **d.** The favored neural representations (i.e. relative-scale vs exact-scale) demonstrated better predictions for task performance. *significant at level 0.05, ** significant at level 0.01.

## Discussion

In the present study, we introduced a novel orthogonal decomposition approach DeCoP that demonstrated superior performance superior to the traditional ‘non-orthogonal’ method in terms of both lower false inference and greater robustness for unbalanced study designs. Further, using DeCoP, we successfully disentangled two functionally independent processes (i.e., evaluation and readiness) from a complex neurobehavioral signal compound during motivational processing (i.e., reward/punishment anticipation during the monetary incentive delay task).

Our findings provide insight into the common ambiguous observations in fMRI tasks that involve multiple interferential latent behavioral or cognitive processes. For example, vmPFC, as a key node in the neural circuitry underlying reward processing and value-based decision making [28–30], was paradoxically ‘inactive’ during the large-win vs neutral contrast. This unexpected ‘inactivation’ could now be understood as a product of a trade-off between two independent processes: activation by reward stimuli (i.e., of the evaluation process) and deactivation as part of the default mode network (i.e., of the readiness process). Similarly, the unbalanced sensitivity toward reward and punishment in the ventral striatum could also be explained as a combination of two balanced independent processes for evaluation and readiness. Furthermore, our findings also demonstrated that the independence of evaluation and readiness processing in the brain putatively is modulated by differential neural circuits targeting VTA and SNc, respectively. This finding provides novel evidence that reward processing is linked to the midbrain dopaminergic system and that evaluation and readiness processes involve distinct underlying neural mechanisms [7, 23, 31].

Further, our novel approach also enables direct statistical inference concerning the functional independence among decomposed latent neurobehavioral processes based on predefined orthogonal latent contrasts. This demonstration may have revolutionary implications for analyzing experimental designs of theoretical neurobehavioral models. While existing research frameworks attempted to identify spatially separated brain regions that activate specifically under a particular neurobehavioral model [5, 18], such a ‘specific’ activation is highly subjective to the selected threshold for significance and hence may not be truly ‘specific’. More problematically, even if the activation is indeed specific to a particular model and null in the other, their underlying signals might still be related. For instance, while the N-shape model was barely activated brain-wide, it nevertheless showed strong negative correlations with the evaluation model in voxels that were ‘specifically’ activated for the evaluation models. Hence, both signals were not independent of each other. Therefore, the statistical framework provided by DeCoP is vital for any meaningful inference for the independence of decomposed signals. Moreover, such a statistical framework enables a computational decomposition for any potentially independent cognitive processes, so long as the experimental design allows a meaningful orthogonal decomposition (i.e. based on the theoretical neurobehavioral models). We therefore expect this new approach to promote new study designs for cognitive processes that were previously theoretically distinct but almost inseparable with common experimental designs.

Finally, DeCoP could comprehensively describe any possible outcomes across all experimental conditions. Complementary to the proposed theoretical neurobehavioral models (for instance the evaluation and the readiness), the other orthogonal components from the same orthogonal basis could also provide valuable information. For instance, in the present study, the complementary orthogonal components, the N-shape and W-shape models, were highly correlated with the evaluation and readiness components, respectively. Therefore, both complementary models modified the response to small monetary stimuli of the corresponding primary models during reward/punishment anticipation. It turns out that while the dACC encoded the exact monetary magnitude of the experimental design in both evaluation and readiness models, most other regions encode only highly abstracted information. These results were consistent with the role of dACC in updating and maintaining subjective value information [32, 33]. Besides dACC, the bilateral insula also favours the exact monetary magnitude scale for the readiness-related signals. However, an unexpected increase in the neutral condition was observed for bilateral insula, rendering the best model in fitness as the W-shape model, instead of the V-shape readiness model. In fact, the W-shape activation was also observed in several previous studies on reward-related decision making [3, 17], though without further discussion. The insula was seemingly involved in two competing processes in regulating incentive salience and the adaptation of external stimulus [34–36], in that higher engagement of dorsal insula could lead to more incentive motivation, but the hyperactive ventral insula was associated with the status of anxiety and maladaptive performance. This observation may help to understand the W-shape activation of aINS as a combination of two competing processes (for instance, positive and negative V-shape activations). Hence, DeCoP could further strengthen our understanding of latent neurobehavioral processes through additionally retrieving complementary components.

## Conclusions

In summary, we have developed and evaluated a universally applicable, novel signal decomposition strategy, ‘DeCoP’, to dissociate behavioral processes that confound the observation of functional neuroimaging signals. This new approach demonstrated superior performance superior to the traditional ‘non-orthogonal’ method in terms of both fewer false inference and higher robustness. Through DeCoP, we demonstrated the independence of evaluation and readiness processing in the brain, putatively modulated differentially by neural circuits targeting VTA and SNc; we also demonstrated that most brain regions, including the ventral striatum, encode signals based on abstract information instead of the observed exact monetary magnitude scale, except for the salience network, i.e., pgACC/dACC and aINS. Most importantly, we demonstrated that DeCoP could help to resolve common paradoxical observations in fMRI tasks which involve complex latent behavioral or cognitive processes, for example, the unexpectedly ‘inactive’ vmPFC in the contrast of large reward vs no reward. We expect that DeCoP could be usefully applied to many other comparably ambiguous datasets and also improve experimental designs for complex cognitive processing.

## Materials and methods

### Participants

The dataset used for this study was selected (see Supplementary Methods for details) from Annual Curated Data Release 2.01 (https://data-archive.nimh.nih.gov/abcd/) of the Adolescent Brain Cognitive Development (ABCD) cohort, which recruited 11,875 children between 9–11 years of age from 21 sites across the United States [20]. The study conforms to each site’s Institutional Review Board’s rules and procedures, and all participants provide informed consent (parents) or informed assent (children). More details of the subjects and the data collection are provided at the ABCD website (https://abcdstudy.org/scientists/protocols/).

### Orthogonally Decoding multi-Cognitive Processes (DeCoP)

Here, we propose a novel approach to decompose each participant’s brain activations at varied conditions (denoted as *y*) with a set of orthogonal basis **x** = (*x_i_*,…*x_k_*), where ‘orthogonal’ means that any pairwise covariances of vectors all equal zero, i.e., *Cov*(*x_i_, x_j_*) = *E*(*x_t_, x_j_*)– *E*(*x_i_*)*E*(*x_j_*) = 0. In this way, the regression coefficients **β** = (*β,…β_k_*) (i.e., the strength of signals) estimated from a multiple linear model with all vectors were the same as those estimated univariately (of simple linear models), i.e., *T*(*β_i_* | *y*, **x**, **β***−i*) = *T*(*β* | *y, x_i_*), where *T*(·) stands for the best linear unbiased estimator. With the above orthogonal settings, the estimated signal strength does not depend on the rest coexisting latent processes and hence is free from signal admixtures. We also proposed that the above individual-level orthogonal decomposition eliminates spurious correlations of signal components (i.e., *β*) introduced by related contrasts (i.e., *x_i_* are correlated), thus allowing us to make meaningful inferences regarding signal independence at the group-level (also see Supplementary Methods).

### The simulation with the real data

We first assumed that the neural response (i.e., the simulated BOLD signal) was indeed a combination of the above two independent signals of evaluation and readiness. Then we generated the simulated BOLD signals with the combinations of the real model-modulated series from 1000 participants randomly sampled from all or part of samples, i.e., *Y* = *β*_1_*X_evaluation_* + *β*_2_*X_readiness_* + *Noise*, where *X_evaluation_* and *X_readiness_* represented the evaluation and readiness model-modulated signals, respectively, *β*_1_ = *β*_2_ ~ *N*(0,1) and *Noise* was the white noise with a variance of 1. Next, we decomposed the simulated signals by the univariate decoding approach (non-orthogonal decoding) and DeCoP and then estimated the differences between the decomposed signals and ground truth. Similarly, we could also generate the simulations under different task designs by randomly selecting the task trials. Each simulation was repeated 1000 times.

We further investigated whether this kind of orthogonal decoding could improve the statistical power of the associations for function-specific phenotypes. We extracted each individual’s brain activation of both evaluation and readiness models decoded by different approaches in NAcc (which responded to both processes simultaneously). For the null hypothesis, we assumed that the simulated phenotype was only associated with evaluation signals. For the alternative hypothesis, we assumed that the simulated phenotype was associated with both evaluation and readiness signals. Then we computed the correlations between the simulated phenotypes and the decoded readiness signals with or without the regression of the corresponding evaluation signals. For each simulation, we simulated with 1000 independent individuals for 1000 times.

## Supplementary data

Supplementary materials consist of a text with Supplementary Methods and Supplementary Results, Supplementary Figures 1-6 and Suppelementary Tables 1-8.

## Author Contributions

Conception or Design of the Study: C.X., T.J., T.W.R. and J.F.. Manuscript Writing and Editing: S.X. and T.J. wrote the manuscript; T.W.R. and J.F. edited the first draft; all authors critically reviewed the manuscript. Imaging Data Preprocessing: S.X., C.X. and W.C.. Visualization: S.X., C.X. and T.J.. Data Collection and Analysis: S.X., C.X., Z.Z. and G.S. conducted all the statistical analyses, under the instruction of T.J. and J.F.. Interpretation: T.J., T.W.R. and J.F.. Supervision of the Study: T.J. and J.F.. Funding Acquisition: T.J. and J.F..

